# Interpretable full-epoch multiclass decoding for M/EEG

**DOI:** 10.1101/2023.03.13.532375

**Authors:** Richard Csaky, Mats W.J. van Es, Oiwi Parker Jones, Mark Woolrich

## Abstract

Multivariate pattern analysis (MVPA) of Magnetoencephalography (MEG) and Electroencephalography (EEG) data is a valuable tool for understanding how the brain represents and discriminates between different stimuli. Identifying the spatial and temporal signatures of stimuli is typically a crucial output of these analyses. Such analyses are mainly performed using linear, pairwise, sliding window decoding models. These allow for relative ease of interpretation, e.g. by estimating a time-course of decoding accuracy, but are computationally intensive and can have limited decoding performance. On the other hand, full epoch decoding models, commonly used for brain-computer interface (BCI) applications, can provide better decoding performance. However, they lack methods for interpreting the contributions of spatial and temporal features. In this paper, we propose an approach that combines a multiclass, full epoch decoding model with supervised dimensionality reduction, while still being able to reveal the contributions of spatiotemporal and spectral features using permutation feature importance. We demonstrate the approach on 3 different task MEG datasets using image presentations. Our results demonstrate that this approach consistently achieves higher accuracy than the peak accuracy of a sliding window decoder while estimating the relevant spatiotemporal features in the MEG signal. Finally, we show that our multiclass model can also be used for pairwise decoding, eliminating the computational burden of training separate models for each pairwise combination of stimuli.

## 1 Introduction

Decoding external stimuli from neuroimaging data, such as Magnetoencephalography (MEG) and Electroencephalography (EEG), has gained increasing attention in recent years (Kay et al., 2008; Cichy et al., 2014). Decoding studies tend to prioritise increasing the discriminatory power (accuracy) between stimuli, e.g. in brain-computer interface (BCI) applications (Koizumi et al., 2018; Cooney et al., 2019a; Défossez et al., 2022), or gaining interpretable insights as to where and when stimuli are represented in the brain (Cichy et al., 2014, 2016). These latter approaches are often referred to as multivariate pattern analysis (MVPA), and typically make use of linear, sliding-window decoders. This allows for the extraction of the interpretable spatiotemporal features that drive the decoding; for example, allowing for the estimation of a decoding accuracy time course (Cichy et al., 2014, 2016; Cichy and Pantazis, 2017; Lappe et al., 2013; Higgins et al., 2022b,a). However, it has been demonstrated that, as one would expect, discriminatory power is also important for the effectiveness of MVPA (Guggenmos et al., 2018). Hence, there is a need in MVPA for decoding methods that improve decoding performance, while maintaining the ability to reveal the spatiotemporal features that underlie the decoding.

One possibility for increasing decoding performance is to abandon the use of sliding window approaches and instead use full epoch decoding. Decoding full-epoch trials has been explored most typically within the context of potential brain-computer interface (BCI) applications, for example in language tasks (Koizumi et al., 2018; Cooney et al., 2019a,b; Hultén et al., 2021; Dash et al., 2020a; Défossez et al., 2022) and motor tasks (Schirrmeister et al., 2017; Dash et al., 2020b; Elango et al., 2017). In contrast with the decoding employed in MVPA, BCI applications often use nonlinear multiclass models (Lawhern et al., 2018). These will generally have good discriminatory power (accuracy), but this comes at the expense of poor interpretability, and are thus not directly useful for MVPA.

Some promising approaches have been investigated recently to make full-epoch models more interpretable, such as the linear forward transform (Haufe et al., 2014). However, this approach can only be applied to linear models. Another option is to apply full-epoch and sliding window decoding on the same data in order to get both perspectives, e.g. in (Ling et al., 2019). Nonetheless, it would be hugely beneficial if a single decoding approach could be used without a loss in performance on both BCI and MVPA.

An additional consideration is computational efficiency. MVPA of MEG data is commonly performed using pairwise decoding methods, i.e. they decode between just two classes at a time (Cichy et al., 2014, 2016; Cichy and Pantazis, 2017; Higgins et al., 2022b,a). When the number of classes gets large, this becomes computationally burdensome. Here, we propose to overcome this through the use of multi-class decoding.

Taking together the aforementioned issues, we propose an approach that can improve decoding accuracy through the use of full-epoch multi-class decoding, while still being able to reveal the underlying spatiotemporal features that drive the decoding. This allows us to consider and investigate the use of neural network decoding models, and we also show the benefit of using supervised feature reduction. We limit our investigations to linear models, leaving nonlinear models for future work. Importantly, to allow access to interpretable features, we make use of permutation feature importance (PFI). PFI is a general technique which can be used to assess which parts of the input contribute the most to the predictions of any black-box model (Altmann et al., 2010). Chehab et al. (2021) have demonstrated the effectiveness of PFI in analysing how certain language features like word frequency affect the forecasting performance of MEG data at various temporal and spatial locations, leveraging a trained encoding model. Deep learning-specific interpretation methods have also been proposed in the context of M/EEG decoding (Schirrmeister et al., 2017; Lawhern et al., 2018)

We assess the proposed approach by systematically comparing it with sliding window decoding on three MEG datasets with visual tasks, finding that our full-epoch decoding outperforms sliding window decoding in terms of accuracy. We then compare PFI with standard alternatives and find that PFI is able to extract the same kind of dynamic temporal, spatial, and spectral information. In addition, we show that pairwise accuracies can easily be gained from a single multiclass model and that these accuracies are on-par with a direct pairwise classification approach.

In short, the aforementioned contributions achieve the best of both worlds: a single decoding model trained on full epochs, empirically good performance, and clear interpretability from an MVPA viewpoint. This approach promises to be useful for both the BCI researcher and the neuroscientist trying to gain insight into the underlying brain activity in a particular task and external stimuli set.

## 2 Material and methods

### 2.1 Data

In this study, we used three visual MEG datasets: two similar datasets from Cichy et al. (2016) and one additional dataset from Liu et al. (2019). The datasets have been collected with appropriate consent from participants and ethical review by Cichy et al. (2016) and Liu et al. (2019), and do not contain any personal information. 15 subjects view 118 and 92 different images, respectively in the first two datasets, with 30 repetitions for each image. The third dataset is part of a larger replay study, and we only use the portion of the data where images are presented in random order for 900ms. Here, 22 subjects view 8 different images, with 20-30 repetitions for each image (depending on the subject). The image sets used in the three datasets are different. We obtained the raw MEG data directly from the authors to run our preprocessing pipeline with MNE-Python (Gramfort et al., 2013). The 118-image and 92-image data are also available publicly in epoched form^1^. We bandpass filtered raw data between 0.1 and 25Hz (50Hz for the 8-image dataset) and downsampled to 100Hz. As recommended by prior work the sampling rate is 4 times higher than the lowpass filer (Higgins et al., 2022b). This is done so that representational alias artefacts are eliminated from the sliding window decoding time courses. We also applied whitening, which involved transforming the data with PCA to remove covariance between channels while retaining all components. The PCA was fit on the training set only but applied to both training and test sets.

In the first two datasets, image presentation lasted for 500ms with an average inter-trial interval of 0.95 seconds. In order to analyse the data using machine learning models, we created two versions of each dataset. The first version consisted of full epochs, with input examples having a shape of [50, 306] (or [90, 273] for the 8-image dataset), where 306 and 273 correspond to the number of MEG channels and 50 and 90 correspond to the number of time points during image presentation. The second version consisted of sliding windows, with input examples having a shape of [10, 306] (or [10, 273] for the 8-image dataset). In this case, we partitioned each trial into overlapping 100ms time windows between 0 and 1000ms post-stimulus and trained separate models on each time window partition as is normally done in the MVPA literature. As a result, 90 independent sliding window models were trained for each dataset.

As opposed to some previous work using a wavelet transform of the trial as features for sliding window decoding (Higgins et al., 2022b), here we use the raw set of timepoints within the respective 100ms window. This means that we rely more on the decoder to extract relevant frequency information rather than directly providing such information in the input. We did compare our approach with the wavelet features and found the latter to be somewhat inferior (see Inline Supplementary Figure 14).

### 2.2 Neural Network with Supervised Dimensionality Reduction (NN)

The Neural Network (NN) method is a four-layer, fully-connected linear neural network which is only run on the full-epoch dataset (Figure 1). The first layer performed a learnable dimensionality reduction, where the full epoch data of dimensions [time points × channels] was multiplied by a weight matrix of shape [channels × components], with components (80) being less than channels. This process is similar to principal component analysis, but in this case, the dimensionality-reducing weight matrix and the decoding model are trained simultaneously; therefore, the dimensionality reduction is optimized for the classification objective. After the first layer, the data was flattened and three affine transformations were applied in sequence (see Figure 1 for dimensionalities). The final layer had an output dimension equal to the number of classes, and the logits from this layer were passed through a softmax function for classification.

**Figure 1:**
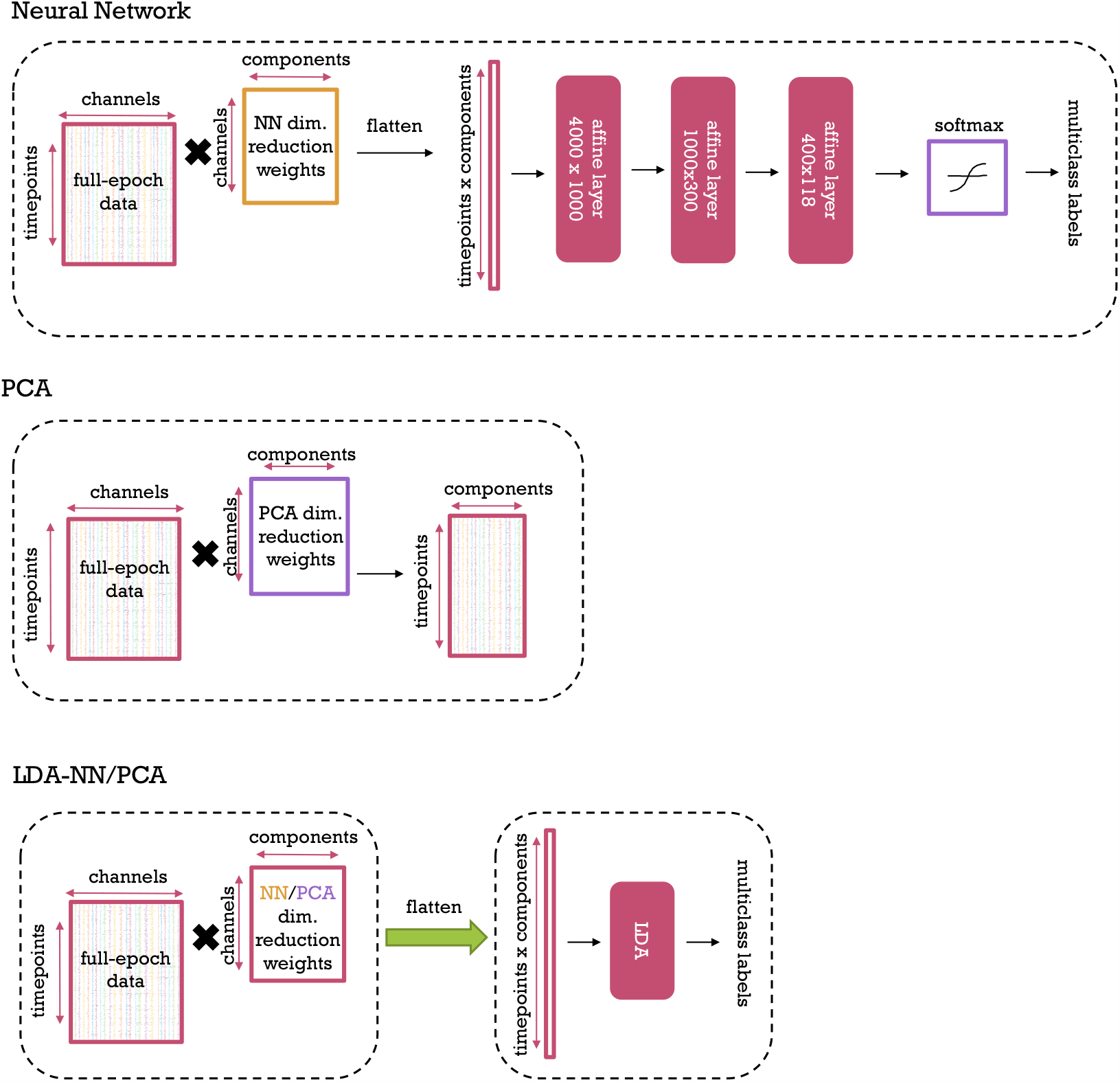
Our Neural Network, PCA, and LDA-NN/PCA methods from top to bottom. Dashed boxes represent separate processing steps, i.e. in the case of LDA-NN and LDA-PCA the respective dimensionality reduction is first used to compute the input features, which are then used to train the LDA model.

The model was trained using cross-entropy loss (Good, 1952) for multiclass classification and included dropout between layers during training (Srivastava et al., 2014). It is worth noting that, as no nonlinearities were used, the model could be replaced with a single affine transformation during evaluation. However, deep linear neural networks are known to have nonlinear gradient descent dynamics that change with each additional layer (Saxe et al., 2013); both the learnable dimensionality-reduction layer and the use of dropout impose additional constraints on the weight matrix during learning.

### 2.3 LDA with Unsupervised Dimensionality Reduction (LDA-PCA)

The LDA-PCA approach has two variants: one that is full-epoch, and one that uses a sliding window. In the full-epoch version, PCA is used to do unsupervised dimensionality reduction on the channel dimension of the full-epoch data as an initial, separate step (Figure 1). The resulting PCA-reduced data matrix, which has a shape of [timepoints x components] is flattened and then used to train a multiclass classifier using LDA.

In the sliding window version, the [timepoints x components] PCA-reduced data matrix is separated into [100ms x 80] windows. The data within each window is then flattened in the same manner as in the full-epoch version and fed into separate LDAs that are distinct to each window.

### 2.4 LDA with Pre-learnt Supervised Dimensionality Reduction (LDA-NN)

In the LDA-NN method, the PCA dimensionality-reducing weight matrix from PCA is replaced with the use of the dimensionality-reducing weight matrix extracted from the pre-trained NN approach (Figure 1). As in LDA-PCA, this weight matrix is then applied to project the data to a [time points x components] shape, after which an LDA model is applied. In the same manner, as LDA-PCA, LDA-NN also has full-epoch and sliding window versions.

### 2.5 Permutation feature importance

To investigate the temporal dynamics of visual information processing, we utilized permutation feature importance (PFI) on our trained models. Specifically, we applied PFI to a trained full-epoch LDA-NN by using sliding windows of 100ms with 1 time point shift for each trial. The information in each window was disrupted by permuting the data across the channel dimension separately for each time window. For instance, if the window was centred around 50ms post-stimulus, the information within that window would be disrupted from 0 to 100ms post-stimulus compared to the original trial, while the rest of the timepoints in the trial remained unchanged. We then evaluated the trained LDA-NN on each of these disrupted trials and compared the accuracy to the original accuracy obtained with the original trials. The greater the accuracy decrease for a trial with disrupted information in a specific time window, the more crucial that time window is to the model’s performance and, therefore, the more information it contains relevant to the model’s objective of discriminating between images. By repeating this analysis for all time windows, we obtain a temporal profile of the information content, similar to the method of training separate models on individual time windows.

Spatial information content can also be obtained in a similar way, but this time by disrupting the information within each channel, by permuting the data across time points for each channel separately. This results in a sensor space map of accuracy decrease, which serves as a measure of visual information content. We compared these sensor space maps to maps obtained by plotting the per-channel accuracy of separate LDA models trained on the full-epoch of each channel. This per-channel LDA method is conceptually similar to using a sliding window in space instead of the traditional time dimension.

We also demonstrated that spatiotemporal information can be obtained jointly using PFI by selecting a window in both space and time (4-channel neighbourhood of 100ms window) simultaneously and disrupting the information within the space-time window, by permuting the values within it over space and time. By sliding this window across all channels and time points, we obtained a spatiotemporal discriminative information content profile.

Finally, we introduce *spectral PFI* to assess the effect of different frequency bands on the visual discrimination objective. First, the data in each channel of each trial is Fourier transformed, and the Fourier coefficients are permuted across channels for each frequency (or frequency band). Then, the inverse Fourier transform is computed, obtaining a trial with disrupted information in specific frequency bands. By applying this method to all frequency bands, we obtained a spectral information content profile, similar to the method of training separate LDA models on features from individual frequency bands (Higgins et al., 2022b). Similar to spatiotemporal PFI we can combine spatial and spectral PFI, by running spectral PFI on a neighbourhood of 4 channels at a time (spatial window) to assess the spectral information content of individual MEG channels. We call this spatio-spectral PFI.

Previous work in our lab applied sliding window decoding in combination with spectral decoding (i.e., training separate models on individual frequency bands), thus assessing the temporo-spectral information content (Higgins et al., 2022b). In order to make comparisons with this work, we developed temporo-spectral PFI. Specifically, after training the full epoch decoding model, we compute the short-time Fourier transform of the entire epoch, using the same parameters as in Higgins et al. (2022b), i.e., a 100ms Hamming window with maximal overlap. We then permuted the channel dimension of one frequency band and one window at a time, leaving the other frequency bands and windows unchanged. Finally, we perform the inverse short-term Fourier transform on the full epoch to get the time domain data back (i.e., channels-by-timesteps), on which the trained decoding model is then applied. By repeating this over all frequency bands and time windows we can obtain the temporo-spectral PFI profile.

### 2.6 Experimental Details

The primary evaluation metric for the three datasets is classification accuracy across the respective number of classes (118, 92, or 8). The main focus of our analysis was on the 118 and 92-image datasets, with the 8-image dataset, included to demonstrate the effects of a much smaller sample size. All of the main results using our decoding methods (NN, LDA-NN, LDA-PCA) are multiclass, with the exception of Figure 9 and the corresponding subsection describing it. In this case for pairwise decoding, we report pairwise accuracy with a chance level of 50%. For all analyses, separate models were fit to separate subjects. Training and validation splits were created in a 4:1 ratio for each subject and class, with classes balanced across the splits. The NN approach was trained for 2000 epochs (full passes of the training data as opposed to epochs in the sense of MEG trials) using the Adam optimiser (Kingma and Ba, 2015). The dimensionality reduction layer and PCA were both set to 80 components, as it is slightly higher than the inherent dimensionality reduction of MaxFilter which is applied to the MEG data, and thus contains more than 99% of variance. We briefly tried our pipeline with 60 components as well on 1 subject and found similar results. The output layer’s dimensionality was equal to the number of classes in the corresponding dataset. Dropout was set to 0.7.

Validation data was not used for early stopping, and the trained NN dimensionality reduction weight matrix (used in LDA-NN) was extracted after the full 2000 epochs of training on the training data. For the LDA models, the shrinkage parameter was set to “auto” using the sklearn package. Comparisons of interest over methods were evaluated using Wilcoxon signed rank tests, with within-subject pairing and subject-level mean accuracies over validation examples as the samples. The PyTorch package was used for training (Paszke et al., 2019), and several other packages were utilized for analysis and visualisation (Pedregosa et al., 2011; Virtanen et al., 2020; Harris et al., 2020; Wes McKinney, 2010; Waskom, 2021; Hunter, 2007). Code written for our analysis can be accessed at https://github.com/ricsinaruto/MEG-transfer-decoding.

## 3 Results

### 3.1 Full-epoch models achieve better accuracy than sliding-window decoding

We set out to test whether full-epoch decoding is better than timepoint-by-timepoint and sliding-window decoding, which are common practices in the M/EEG literature (Carlson et al., 2011, 2013; Su et al., 2012; Ramkumar et al., 2013; Cichy et al., 2017; Grootswagers et al., 2017; Kurth-Nelson et al., 2016; Liu et al., 2019; Higgins et al., 2022b). We first selected the best-performing linear classifiers trained for multiclass decoding from a set of commonly used candidate models, including support vector machines (SVM), linear dis-criminant analysis (LDA), logistic regression, and Lasso. The results, depicted in Figure 2, indicate that LDA performed the best among the examined models in both average and peak classification accuracy, with logistic regression exhibiting comparable performance. For this reason, and as described in the methods, we used LDA in all further analyses for comparing different classification strategies.

**Figure 2:**
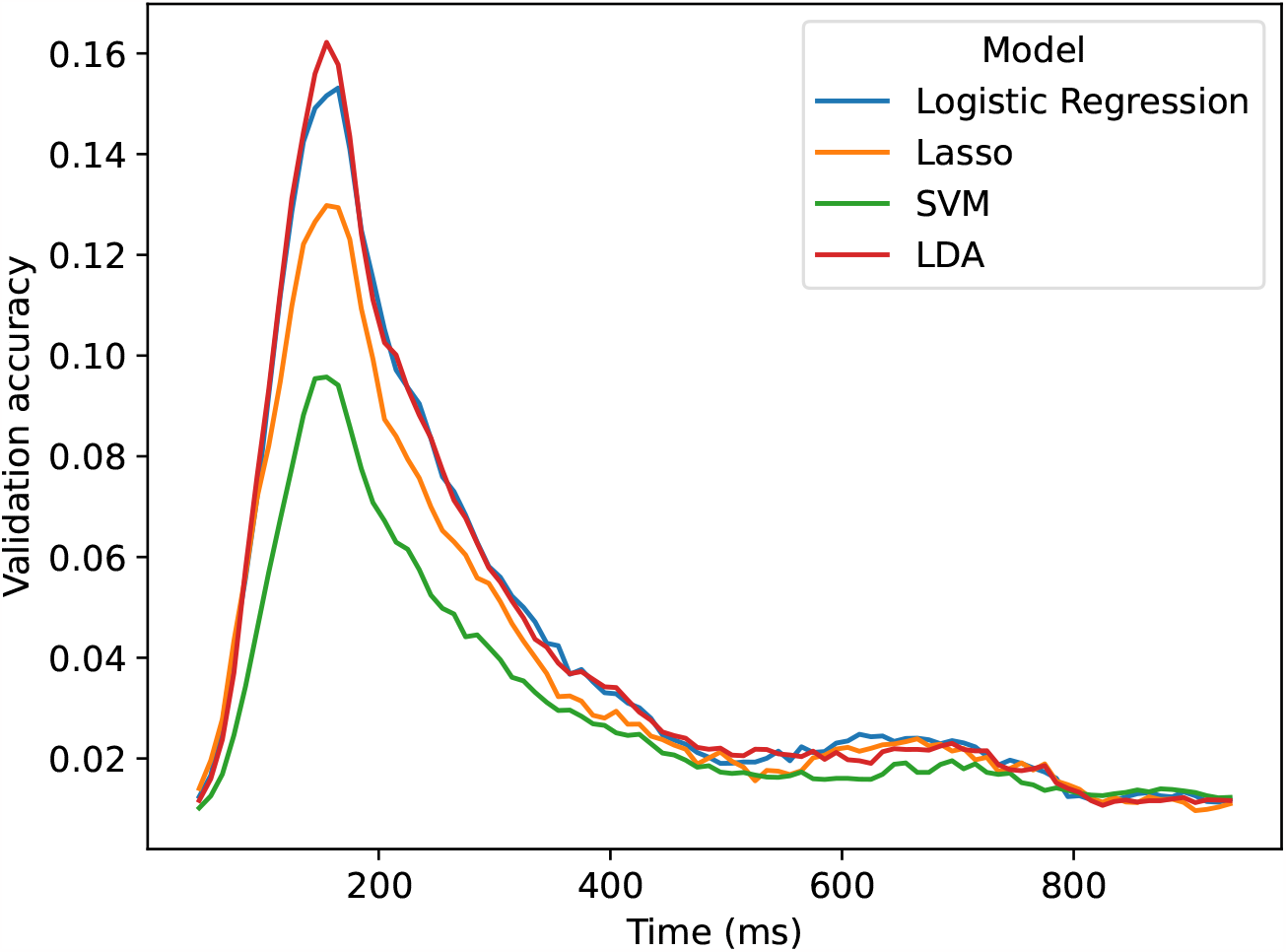
Comparing different sliding window models trained on PCA features on the 118-image dataset for multiclass decoding. The sliding window size is 100ms. Results are averaged across subjects.

The performance of multiclass full-epoch models was compared to that of sliding-window decoding for both LDA-PCA and LDA-NN on the three datasets in Figure 3. The peak performance of sliding-window decoding was observed at 150-160 ms post-stimulus for the 92 and 118-image datasets, and at 200 ms post-stimulus for the 8-image dataset. These findings are broadly consistent with previous research on the temporal dynamics of visual information processing in MEG (Cichy et al., 2014, 2016; Cichy and Pantazis, 2017; Higgins et al., 2022b; Liu et al., 2019; Guggenmos et al., 2018). For the 92 and 118-image datasets a second smaller peak was observed around 650-660 ms post-stimulus. As the image presentation is switched off at exactly 500ms, we reason that the second peak is due to the brain reacting to this event. The first peak is observed 150-160 ms post-stimulus onset, while the second peak occurs 150-160 ms post-stimulus offset.

**Figure 3:**
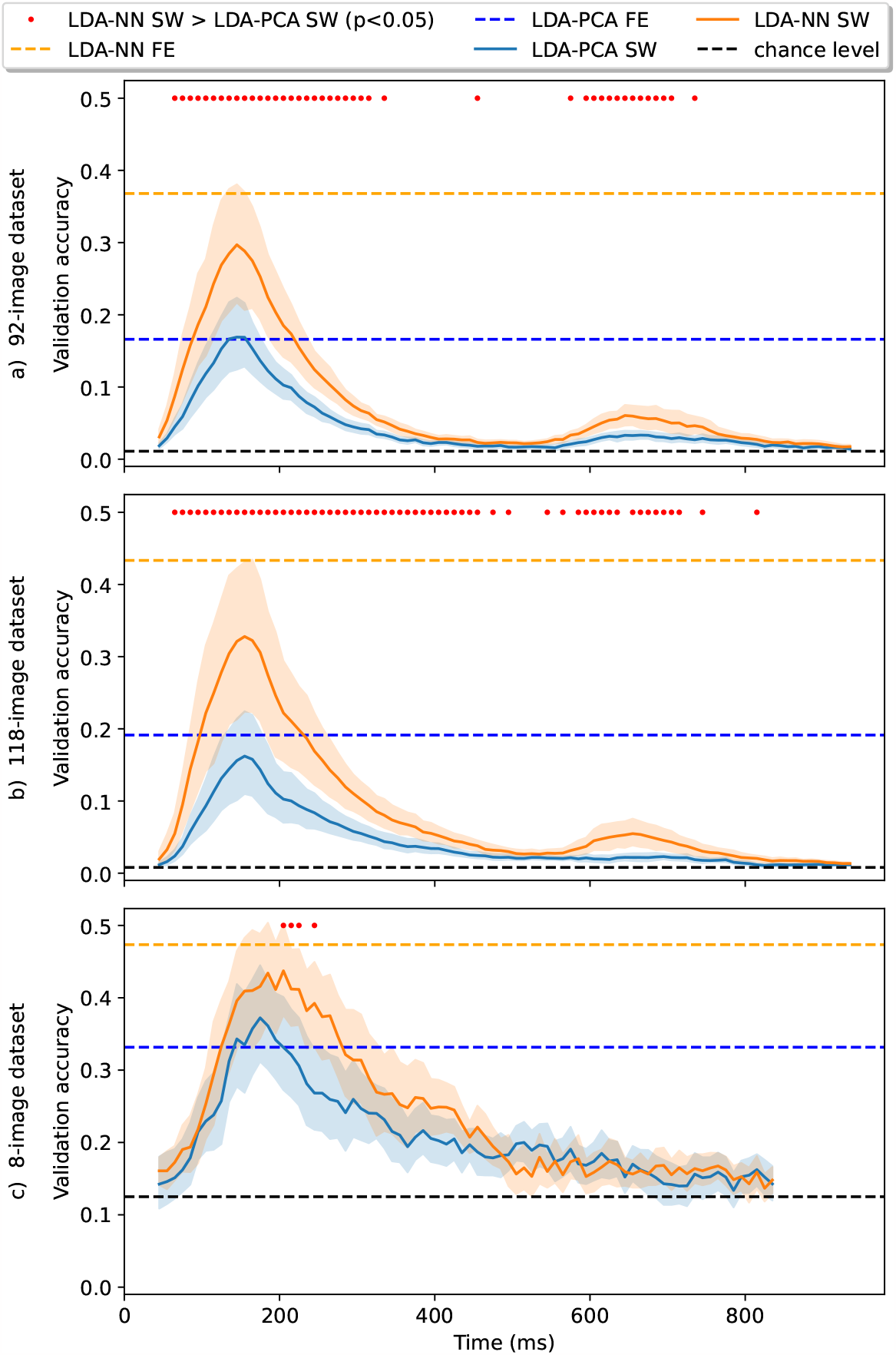
Models trained on the sliding-window versions of the 92-class dataset (top), 118-class dataset (middle) and 8-class dataset (bottom) for multiclass decoding. Wilcoxon signed-rank tests are reported between sliding window LDA-NN and LDA-PCA. FE stands for full-epoch models, and SW stands for sliding window models. The blue and orange dotted lines are placed at the average performance of full-epoch LDA-NN and LDA-PCA, respectively. All statistical tests are corrected for multiple comparisons across all time points (i.e. p-values are multiplied by 90). Shading indicates the 95% confidence interval across subjects. For the full-epoch results, please see Figure 4 for distributions across subjects. LDA-NN is better across almost all time points than LDA-PCA, and full-epoch accuracy is higher than peak sliding window accuracy for both LDA-NN and LDA-PCA (except in the 92-class and 8-class datasets).

Across subjects, the full-epoch LDA-PCA approach demonstrated significantly higher accuracy than the best sliding-window LDA-PCA approach on the 118-class dataset (3.1% increase, p < 1e-4). On the 92-class dataset, no significant difference was observed between these models, though full-epoch LDA-PCA still outperformed the sliding-window version at most time windows. A similar comparison between full-epoch LDA-NN and peak sliding-window performance showed that full-epoch models had higher accuracy on both the 92- and 118-class datasets (7.1% and 10.5% increase, respectively, p < 1e-4). The tests were corrected for multiple comparisons across time points. These results indicate that training a model on the full epoch generally leads to better performance than using the best sliding-window model, except for the LDA-PCA approach on the 92-image dataset. However, as noted in the following section, it is advisable to use an LDA-NN model in any case.

Our results could be affected by the choice of window size for the sliding window LDA (100 ms). Thus, we repeated the sliding window LDA for different window sizes, including a window of 1 sample (i.e., timepoint-by-timepoint decoding), and the results are presented in Inline Supplementary Figure 10. We found that as the window size increased accuracy improved reaching full-epoch performance with a 200ms window but the accuracy profile became more distorted and the peak shifted compared to the results obtained with a single time point. Finally, on the 8-image dataset, the full-epoch model had higher accuracy than the peak sliding-window model, though this difference was not significant. It should be noted that the reduced effectiveness of the full-epoch model on this dataset may be due to both the longer epoch of 900 ms and the smaller amount of data. This can lead to overfitting due to a larger number of features and fewer examples.

### 3.2 Supervised dimensionality reduction is better than PCA

We next investigated the effect of incorporating a learned, supervised dimensionality reduction layer in our models, i.e. a dimensionality reduction optimised to aid a downstream classification task. We, therefore, modified the LDA-PCA approach by replacing the unsupervised dimensionality reduction performed by PCA with the supervised dimensionality reduction (of equal dimensionality) from the Neural Network (NN) approach, as described in Section 2. We refer to this modified approach as LDA-NN. As shown in Figure 4, this simple change resulted in a significant improvement in performance (20.2% for the 92-class dataset and 24.2% for the 118-class dataset, p < 1e-4). We also assessed the performance of the pure NN model and found that it has a similar performance to LDA-NN. In other words, the supervised dimensionality reduction effectively eliminated the performance gap between the LDA and the Neural Network (NN) approach.

**Figure 4:**
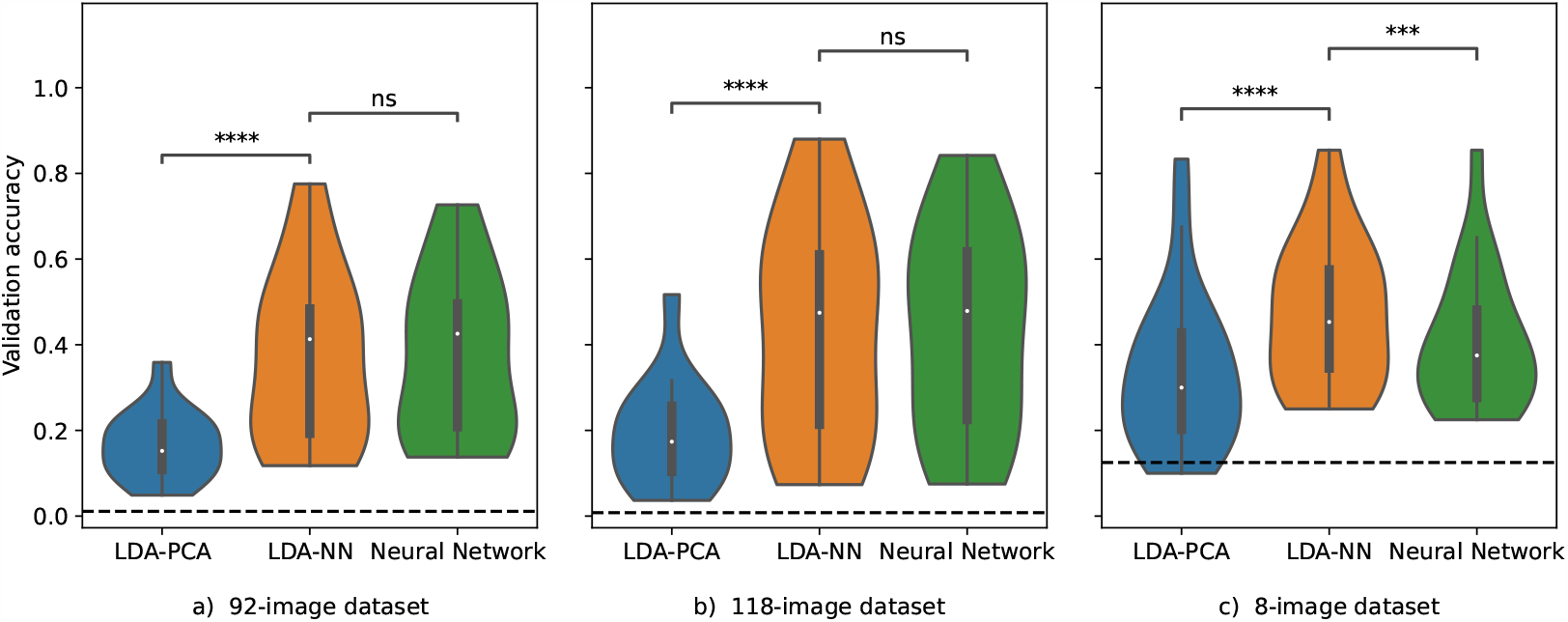
Models trained on the full-epoch versions of the 92-class (left), 118-class (middle), and 8-class (right) datasets for multiclass decoding. The violin plot distributions are shown over the mean individual subject performances. The dashed black line represents the chance level. Wilcoxon signed-rank tests are shown where 4 stars mean p < 1e-4, and 3 stars mean p < 1e-3. “ns” means that the p-value is higher than 0.05.

The sliding window versions of LDA-PCA and LDA-NN are also compared in Figure 3. Across most time points (and all time points around the 2 peaks), LDA-NN is significantly better than LDA-PCA, when corrected for multiple comparisons across time points. Similar conclusions can be drawn on the 8-image dataset, although LDA-NN is better than the NN approach, possibly due to the reduced performance of neural networks on small datasets in general. In summary, our results suggest that using a full-epoch LDA-NN or a simple linear Neural Network results in the best performance across all datasets and that the feature reduction should be learned in a supervised manner for both the LDA and Neural Network models.

### 3.3 Full-epoch models contain the same kind of temporal and spatial information as sliding window decoding

One of the benefits of sliding window or time-point-by-time-point decoding is that it is straightforward to obtain a time course of decoding accuracy (e.g., Figure 3), allowing for interpretation of the temporal dynamics of neural representations. Here we show that full epoch decoding in combination with permutation feature importance (PFI) can give the same qualitative information. The results presented in Figure 5 indicate that temporal PFI applied to a full-epoch LDA-NN model produces temporal profiles similar to those obtained using sliding window LDA-NN models with a window size of 100ms across all three datasets. The peak sliding window performance also aligns well with the peak accuracy loss for PFI.

**Figure 5:**
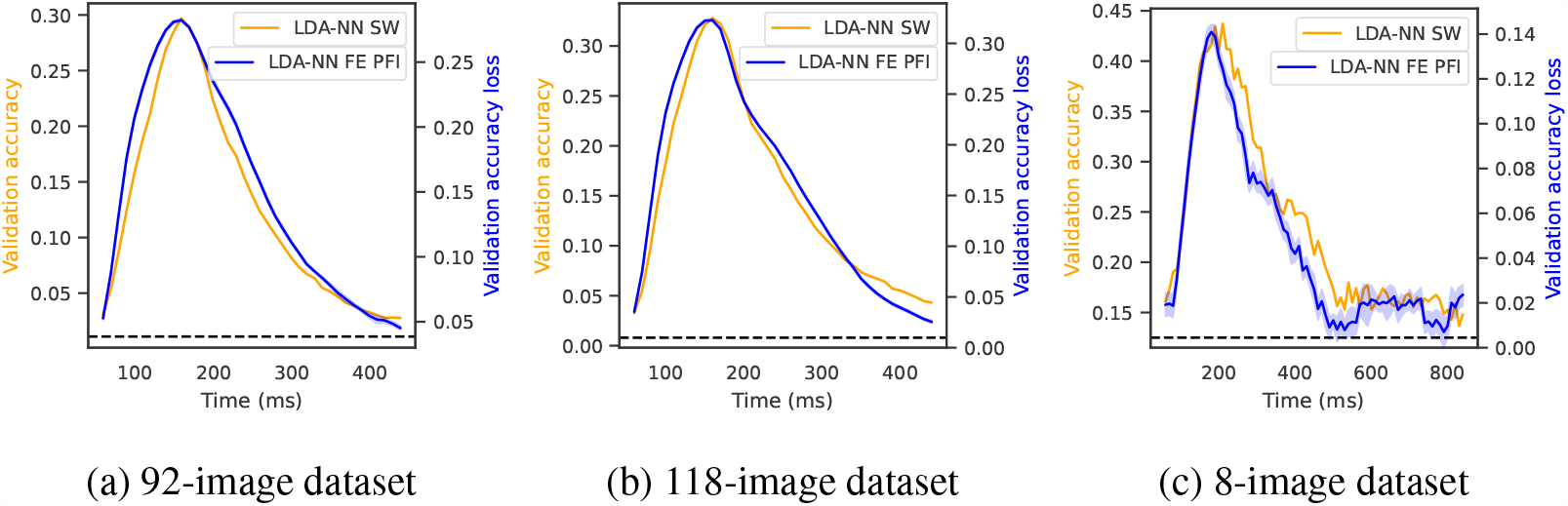
Comparison of multiclass sliding window LDA-NN (orange) and the temporal PFI of multiclass full-epoch LDA-NN (blue) across the three datasets. Results are averaged across all subjects in the respective datasets, and shading indicates 95% confidence interval across permutations for PFI.

In addition, we investigated the ability of PFI to accurately capture spatial information by applying it to a full-epoch LDA-NN model on the 118-image dataset. To do this, we permuted time points from the gradiometers and magnetometers located at the same position in the MEG data simultaneously to obtain a single sensor space map. We compared these to the maps obtained by training separate LDA models on the full epoch of the same three sensors (2 gradiometers and 1 magnetometer). This approach can be viewed as a sliding window across space. All PFI results are averaged over the accuracy losses of individual subjects, which can somewhat smear both spatial and temporal profiles. The results, shown in Figure 6, demonstrate good alignment between the accuracy loss of spatial PFI and persensor accuracy of LDA-NN, indicating that PFI can effectively recover spatial information content.

**Figure 6:**
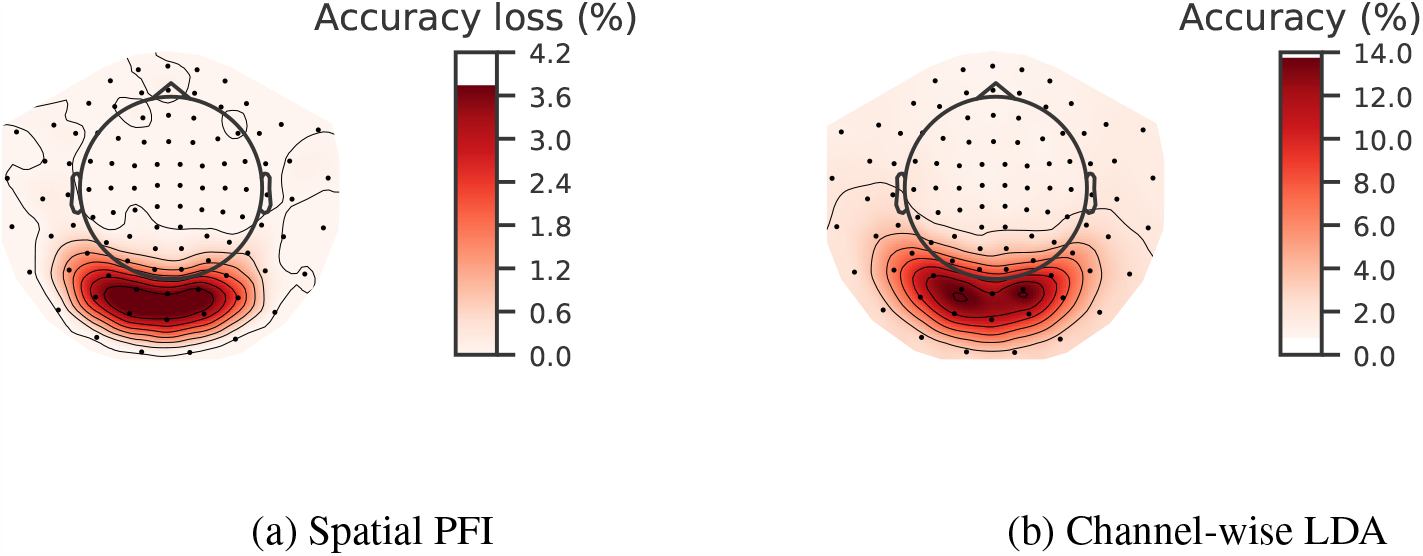
Comparison of multiclass channel-wise LDA model (b) with the spatial PFI of multiclass full-epoch LDA-NN (a). Spatial maps are averaged across all 15 subjects on the 118-image dataset. Both PFI and the channel-wise LDA model are run on 3-channels in the same location at a time (1 magnetometer and 2 gradiometers).

We also employed PFI to extract spatiotemporal information jointly from a trained full-epoch LDA-NN model on the 118-image dataset. Specifically, we used a 100 ms time window and a 4-channel spatial window (i.e., the 2 gradiometers and 1 magnetometer on three sides of the sensors in question) for each time point and channel, shuffling the values within these blocks. This allowed us to unravel the temporal and spatial information simultaneously, showing that only channels located in the visual area exhibited the characteristic temporal profile and that there was a gradient with channels further from the visual area displaying progressively lower peak accuracy loss (Figure 7). Additionally, we observed that the temporal evolution of the sensor space maps showed the visual area sensors to be consistently the most important for the decoding objective across all time points. A full animation of the temporal evolution of the sensor space maps is provided in Inline Supplementary Video 1. In theory, the sliding window LDA and the per-channel LDA approach could be combined to get a similar spatiotemporal profile, where each LDA model is trained on the sliding window of 4 channels at a time. However, in practice accuracy might suffer substantially with so few input features, and it would be computationally taxing considering the amount of LDA models required to train. Overall, PFI proved to be a useful technique for investigating full-epoch data and obtaining spatiotemporal information similar to what can be obtained from individual sliding window models.

**Figure 7:**
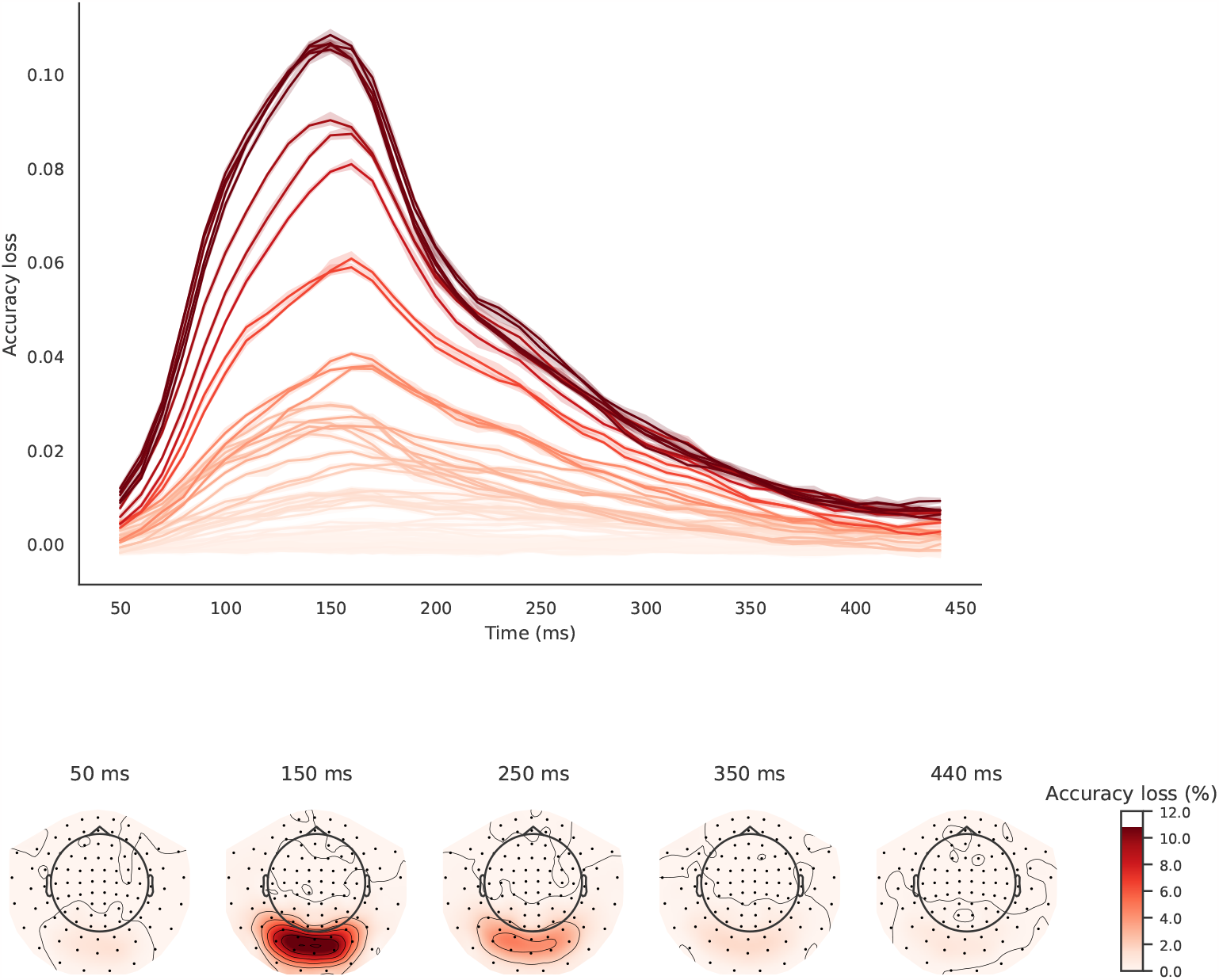
Spatiotemporal PFI of multiclass full-epoch LDA-NN on the 118-image dataset. Blocks of 4-channel neighbourhoods and 100ms time windows are shuffled to obtain a spatial and temporal profile jointly. Each line in the temporal profile corresponds to a sensor, and each sensor space map is obtained with a time window centred around the respective time point. The shading in the upper plot is across the 10 permutations used for PFI and indicates the 95% confidence interval. Both temporal and spatial profiles are averaged over subjects.

Finally, Figure 8a presents our spectral PFI results averaged over subjects. Because of the sampling rate of the data and the size of the epochs, the frequency resolution is 2Hz. There is a clear peak of spectral information content at 4Hz, after which it rapidly declines, as is expected in MEG data (i.e., the 1/f characteristic). Similar to spatiotemporal PFI we can combine spatial and spectral PFI to assess the spectral information content of individual MEG channels (see Inline Supplementary Figure 13).

**Figure 8:**
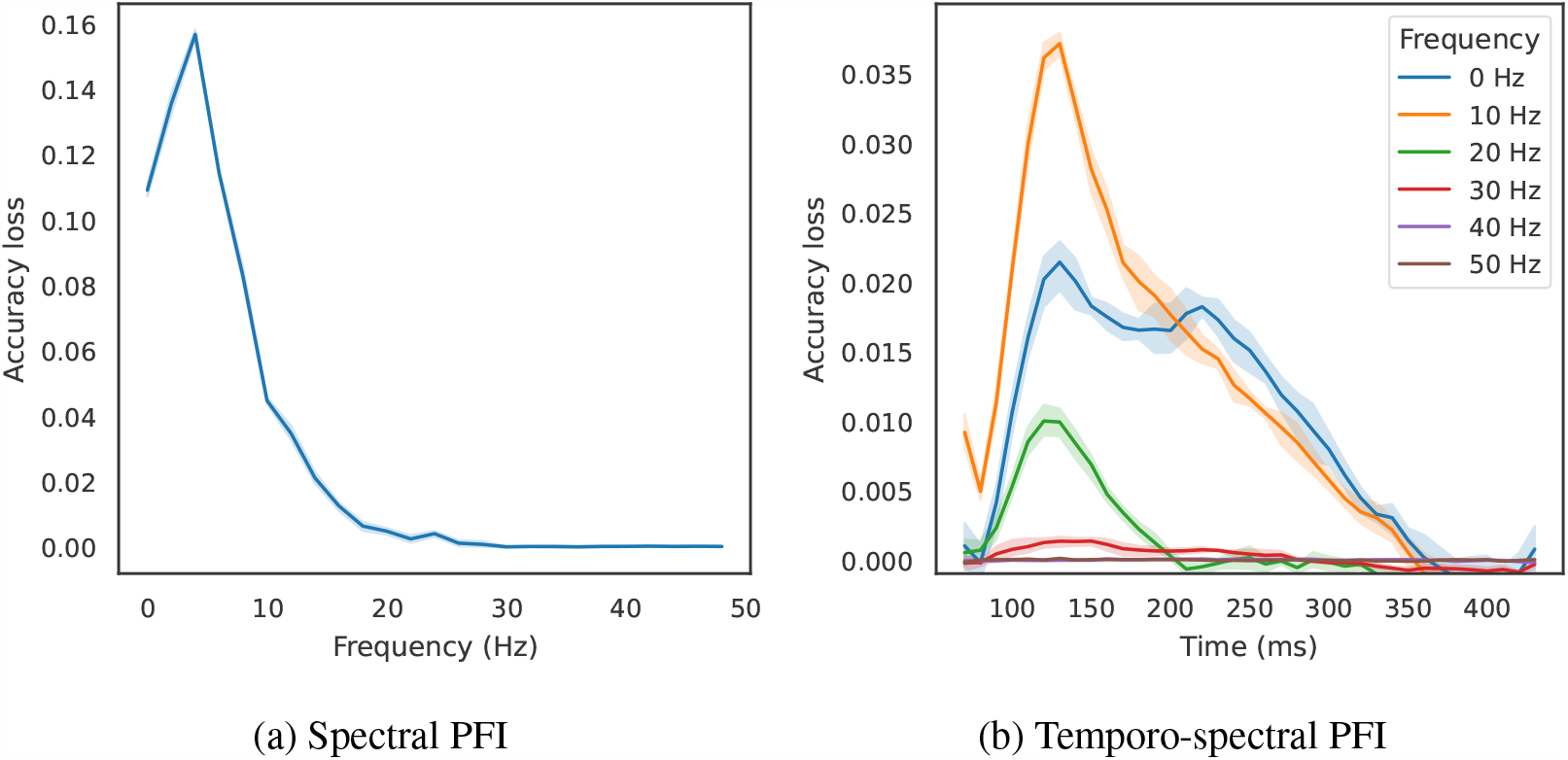
Spectral PFI (left) and temporospectral PFI (right) of multiclass full-epoch LDA-NN on the 118-image dataset. Shading indicates 95% confidence interval across permutations. Results are averaged across subjects.

We present temporospectral PFI in Figure 8b, an alternative method to showing tempo-ral information content within individual frequency bands using separate LDA models trained on wavelet features (Higgins et al., 2022b). When permuting a specific time window, we also permuted the frequency content of the time window right before and after, to obtain a smoother temporal profile. As expected from the standard temporal PFI, the temporal peak is between 100 and 150ms. Spectrally, the 10Hz band is the most important to the decoding objective, with higher bands being less and less useful, confirming the observations in Higgins et al. (2022b). Note that in later timepoints (>200ms) the 0Hz band seems to be slightly more important than the 10Hz band. We hypothesise that because of the reduced frequency resolution of the short-time Fourier transform, the 4Hz peak observed in spectral PFI gets represented in the 10Hz band in temporo-spectral PFI, and that the actual peak would be 4Hz if using a higher resolution.

### 3.4 Multiclass decoding models evaluated for pairwise decoding are on par with direct pairwise models

So far, we have demonstrated that multiclass full-epoch models are better than sliding window models while maintaining the same level of spatiotemporal information. Here we wish to highlight an additional advantage of using multiclass full-epoch models. Researchers frequently use pairwise models to analyse the representational differences between individual conditions or groups of conditions, such as in representational similarity analysis (RSA). However, this approach can be computationally intensive, especially when dealing with a large number of classes.

Here, we show how we can utilise a single trained multiclass full-epoch LDA-NN model to predict pairwise accuracy scores. This is done by iteratively taking all pairs of conditions, computing the predicted probabilities across all classes for each trial, and selecting the condition with the higher probability (of the two conditions) as the predicted class. By comparing this to the ground-truth labels, we can obtain pairwise accuracy scores for each pair of conditions. In Figure 9, we compared the results of this method with those obtained by training individual pairwise (full epoch LDA-NN) models as is typical in the literature. For the 92 and 118-image datasets, the multiclass model achieved slightly, but significantly higher pairwise accuracy than the individual pairwise models. The difference was not significant for the 8-image datasets. Therefore, using a multiclass model can yield pairwise results that are similar to or even better than those obtained from individual pairwise models. This provides a much more efficient way of obtaining pairwise accuracies for the purposes of RSA.

**Figure 9:**
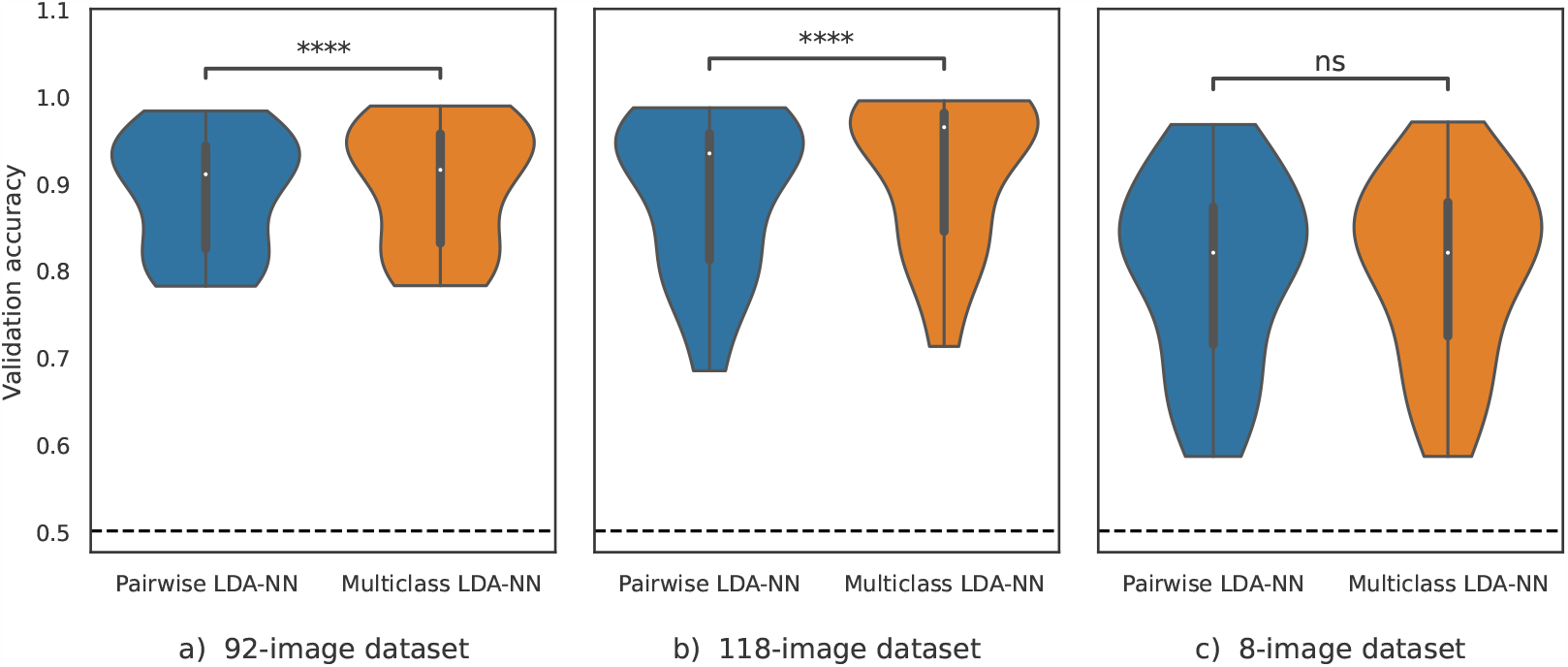
Comparison of pairwise full-epoch LDA-NN models (blue) with multiclass models evaluated for pairwise classification (orange) across the three datasets. In all datasets except the 8-image dataset, multiclass models evaluated in a pairwise fashion are significantly better (****, p<1e-4). The violin plot distributions are shown over the mean individual subject performance. The dashed line represents chance level.

## 4 Discussion

We made the following contributions in this work. We showed empirically that full-epoch models achieve higher accuracy than sliding window decoding models. We showed how temporal, spatial, and spectral brain activity patterns related to stimulus discrimination can be extracted for any black-box full-epoch model. We showed how pairwise accuracies can be gained from a single multiclass model, and that these are on par with direct pairwise decoding. We proposed to learn the dimensionality reduction of input features in a supervised way, improving decoding performance substantially. Next, we discuss each result in more detail.

We have found that training a single full-epoch model for multiclass decoding is effective in improving decoding performance, and have shown how this can be used while still providing neuroscientific insights by using PFI to learn which features are contributing to the decoding accuracy. Our results show that a full-epoch model generally performs better than individual sliding window models for visual decoding tasks, and the magnitude of this effect increases with the size of the dataset. Even with smaller datasets, such as the 8-image dataset used in our experiments, the time-efficiency benefits of using a full-epoch model are still evident, as training sliding window models takes roughly 10 times longer than a single full-epoch model for a 100ms time window with a 100Hz sampling rate. Additionally, our analysis of different window sizes (see Supplementary Material) showed that while larger window sizes may improve performance, they are not effective in accurately capturing the temporal profile of information content. It has also been suggested that using equal-length time windows for all trials does not account for trial-by-trial variability, and Vidaurre et al. (2018) proposed time-resolved decoding using a Hidden Markov Model to segment trials along the time dimension. This approach still involves training multiple models on multiple time windows. We, therefore, recommend using full-epoch models, as they only need to be trained once and can be used to select any desired window size for temporal or spatial investigations through PFI, providing good decoding performance and dynamic spatiotemporal resolution without the need for retraining.

We also found that incorporating a supervised dimensionality reduction layer is essential for good decoding performance when using linear neural networks and LDA models. This can be used as a drop-in replacement over standard unsupervised dimensionality reduction typically done with PCA. In our approach, we used pre-trained weights from a neural network for the feature reduction part of the LDA-NN model and kept this part fixed during the training of the LDA. While we hypothesise that constructing an LDA-NN model, in which both the feature reduction weights and the LDA model are trained together, would produce similar results, this has not been tested.

We compared PFI results from a full-epoch model with those from individual models trained on either separate time or spatial windows. This demonstrated that PFI can effectively extract both temporal and spatial information, and can also be used to investigate the interaction between these two dimensions. We also introduced a new technique whereby PFI can be used to extract spectral discriminatory information content and confirmed that this matches previous work training individual models on separate frequency bands. PFI is a particularly flexible technique, as it can be applied to nonlinear models and temporal or spatial resolution can be chosen post-hoc without the need for retraining. The performance of full-epoch nonlinear decoding and corresponding PFI analysis will be explored in future work. PFI can also be applied to individual conditions or single trials by rerunning with different permutations, enabling the investigation of various neuroscientific questions. Other methods for obtaining temporal and spatial information from trained models, such as the Haufe transform, are limited to linear models and do not provide trial-level patterns (Haufe et al., 2014). As opposed to the statistical nature of PFI, the Haufe transform directly maps the weights of a linear decoding model to input patterns, thus showing which parts of the input are the most important for the decoding objective. One downside of PFI compared to the Haufe transform is that the absence of influence on the output does not necessarily mean that those parts of the input (channels or time windows) do not contain information about the target.

In addition to the benefits of using a full-epoch multiclass model for MVPA, we have demonstrated a simple method for obtaining pairwise accuracies that are comparable or even superior to those obtained through individual pairwise models. This approach is useful for RSA and reduces computation time by approximately half the number of conditions in the data, as pairwise models reuse this data for training while a multiclass model uses it only once. Although the data must still be reused for evaluation, we can assume that evaluation is much faster than training. The slight increase in performance when using multiclass models could be because decoding many classes together helps to better constrain the relationship between features and class labels compared to doing 2 classes at a time.

To conclude, we recommend using a full-epoch multiclass model equipped with a supervised dimensionality reduction in order to achieve the best possible decoding performance while also allowing for flexibility in conducting neuroscientific investigations post-hoc such as MVPA or RSA. Our methods and recommendations scale well with data size and can be readily applied to deep learning models as well, thus bringing the applications of decoding to brain-computer interfaces and representational brain dynamics under a joint approach.

## Supporting information

Inline Supplementary Video 1

## Acknowledgments

This research was supported by the NIHR Oxford Health Biomedical Research Centre. The views expressed are those of the author(s) and not necessarily those of the NIHR or the Department of Health and Social Care. RC is supported by a Wellcome Centre Integrative Neuroimaging Studentship. MVE’s research is supported by the Wellcome Trust (215573/Z/19/Z). OPJ is supported by the UK MRC (MR/X00757X/1). MWW’s research is supported by the Wellcome Trust (106183/Z/14/Z, 215573/Z/19/Z), the New Therapeutics in Alzheimer’s Diseases (NTAD) study supported by UK MRC and the Dementia Platform UK (RG94383/RG89702) and the EU-project euSNN (MSCA-ITN H2020-860563). The Wellcome Centre for Integrative Neuroimaging is supported by core funding from the Wellcome Trust (203139/Z/16/Z).

## A Supplementary Material

### A.1 Window size comparison

In Figure 10, we examined the impact of the sliding window size on the results of our LDA-NN models. We trained models using sliding window sizes of 10ms, 100ms, 200ms, 300ms, and 400ms. As expected, using a single time point (10ms) resulted in lower accuracy compared to a 100ms window. As the window size increased, we observed two trends. First, accuracy improved and the peak accuracy of a 200ms window already reached the full-epoch level. Second, the accuracy profile became more distorted and the peak shifted compared to the results obtained with a single time point. In some cases, full-epoch performance was even exceeded by a few percentage points with a 300ms window. This may not be surprising, as a larger window that focuses on the most significant part of the input results in fewer features compared to using the full epoch. However, it is advisable to avoid using a window larger than 100ms in sliding window analysis due to its distortion and lower temporal resolution. One potential solution could be to combine the sliding window models with PFI analysis, but this would be inefficient. We can therefore conclude that using a full-epoch model is the optimal solution, even if it results in slightly lower accuracy. Additionally, we expect that with larger datasets, full-epoch models would outperform sliding window models regardless of window size, as the ratio of features to examples would be reduced.

**Figure 10:**
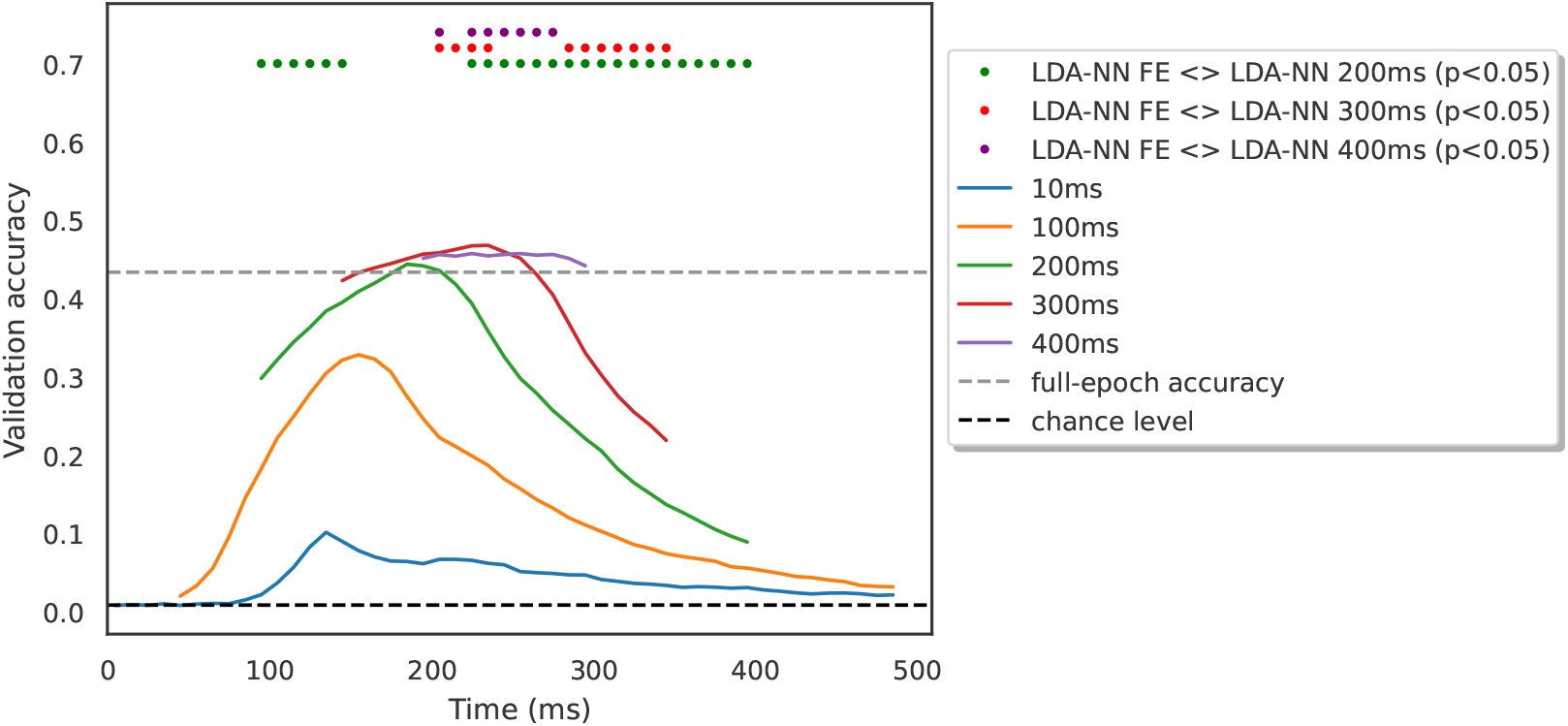
Comparing sliding window LDA-NN with different window sizes on the 118-image dataset. Results are averaged across subjects. Wilcoxon signed-rank tests are reported between the sliding window models and the full-epoch model, corrected for all comparisons in the figure.

### A.2 Inverse temporal and spatial PFI

We investigated an alternative method of performing PFI, referred to as inverse PFI. This method is not common in the literature, but could be interesting from an MVPA viewpoint. Inverse PFI differs from standard PFI in that it shuffles values outside a specified time window, rather than within it. Standard PFI assesses the impact of disrupting information within a specific window on performance and therefore reveals the importance of that window for discriminating between images. In contrast, inverse PFI investigates performance when all information outside a specified window is disrupted, thereby providing insight into the performance that can be achieved using only the information contained within the time window. The temporal PFI results for both standard and inverse PFI are presented in Figure 11. While both approaches are similar to the standard sliding window LDA profile, there are some differences as well. We also conducted this analysis in the spatial domain, the results of which are shown in Figure 12. In this domain, the inverse PFI approach exhibits less contrast between visual channels and other channels but appears to distinguish between visual channels more similarly to channel-wise LDA than standard PFI. It is not the aim of this study to determine which approach is superior, as both seem to have their merits.

**Figure 11:**
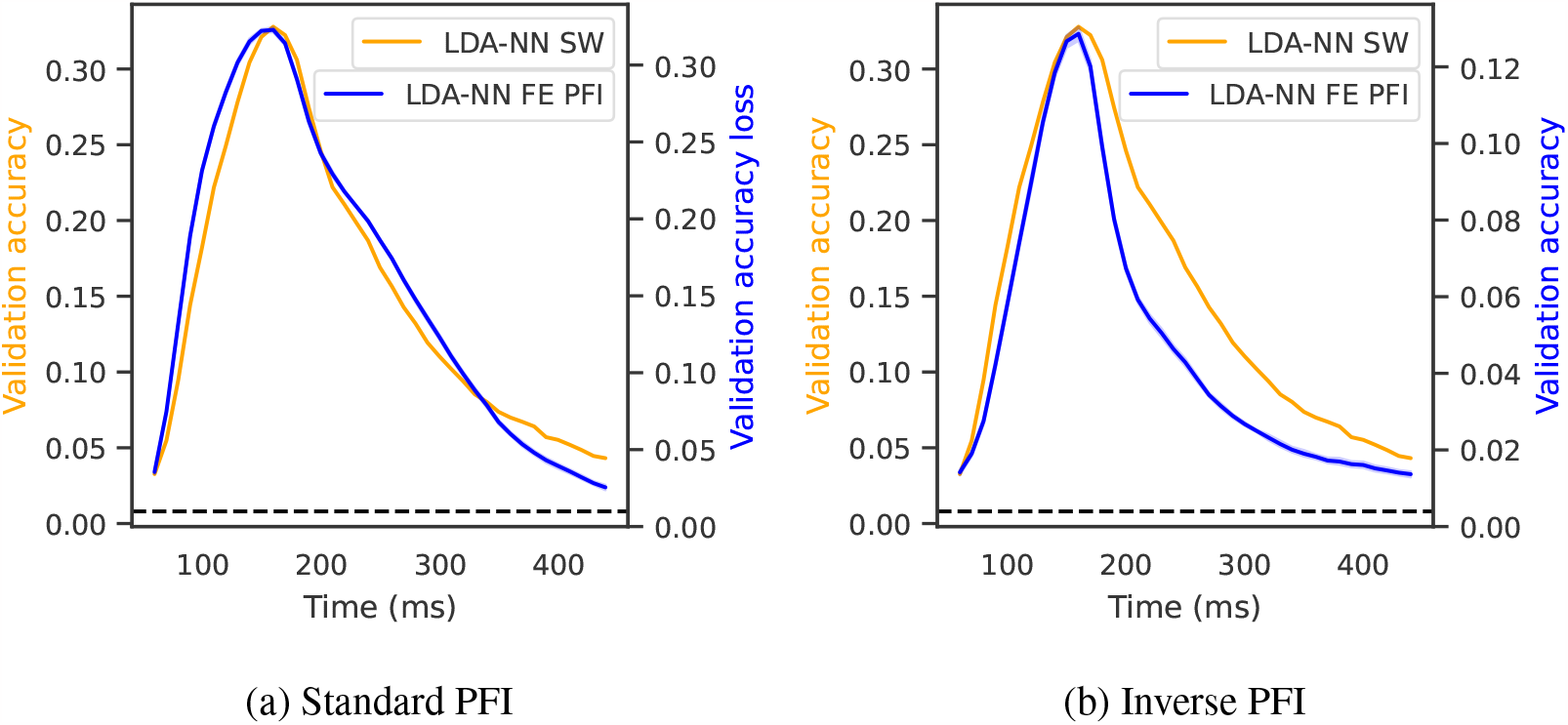
Comparison of multiclass sliding window LDA-NN (orange) with standard temporal PFI (a) and inversed temporal PFI (b) using a trained LDA-NN model on the 118-image dataset. Results are averaged across subjects, and shading indicates the 95% confidence interval across permutations for PFI.

**Figure 12:**
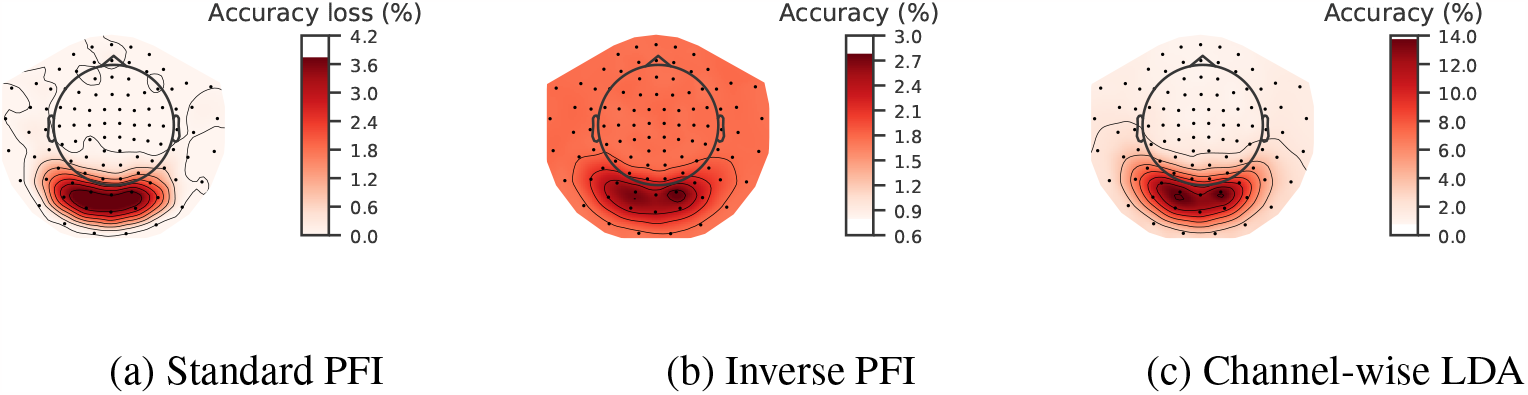
Comparison of channel-wise LDA model (c) with the standard spatial PFI (a) and inverse spatial PFI (b) of full-epoch multiclass LDA-NN. Results are averaged across all 15 subjects on the 118-image dataset. Both PFI and the channel-wise LDA model are run on 3-channels in the same location at a time (1 magnetometer and 2 gradiometers).

**Figure 13:**
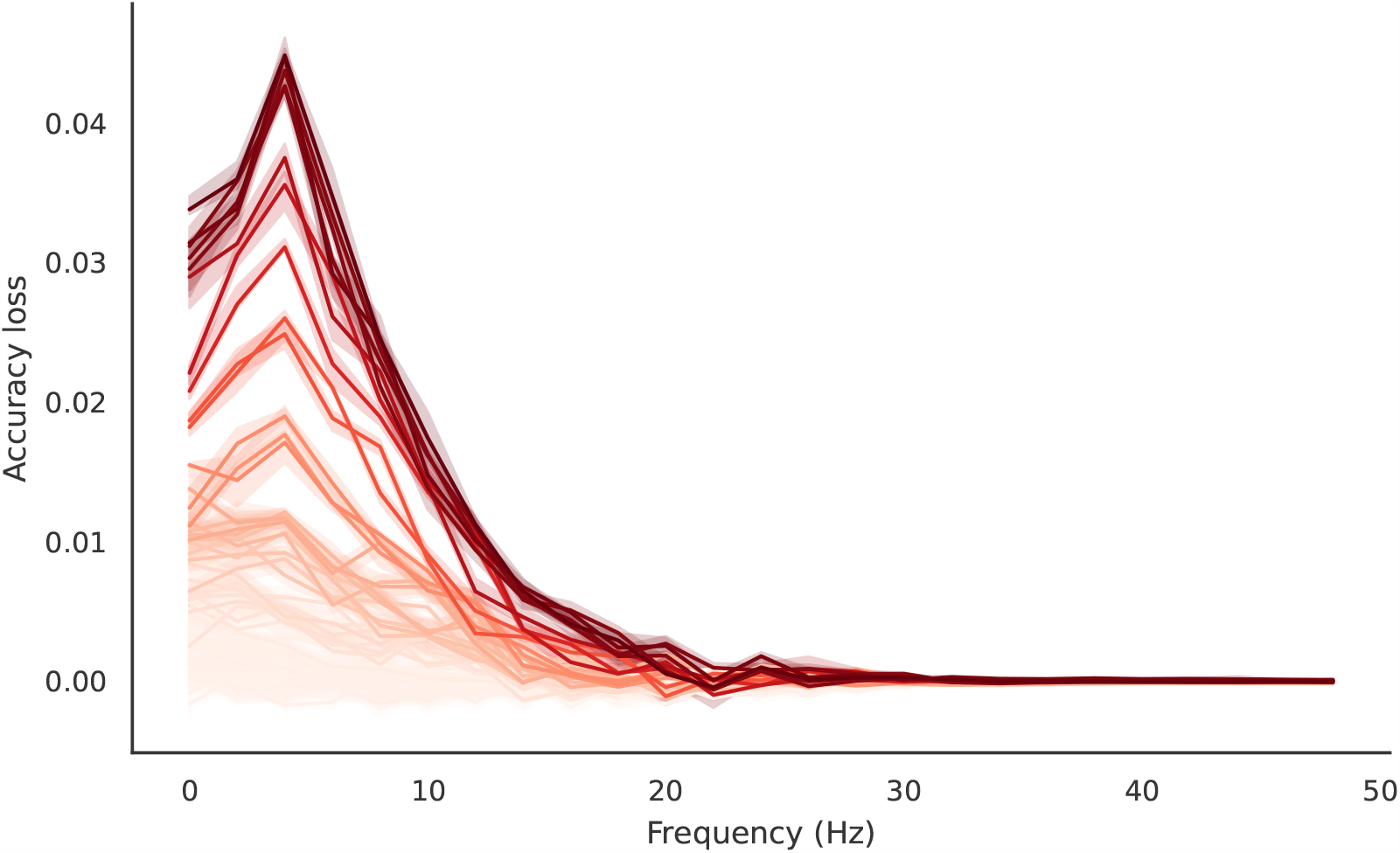
Spatiospectral PFI of multiclass full-epoch LDA-NN on the 118-image dataset, averaged over subjects. Blocks of 4-channel neighbourhoods are shuffled in each frequency to obtain the per-channel frequency profile. Each line corresponds to a sensor. The shading is across the permutations used for PFI and indicates the 95% confidence interval.

**Figure 14:**
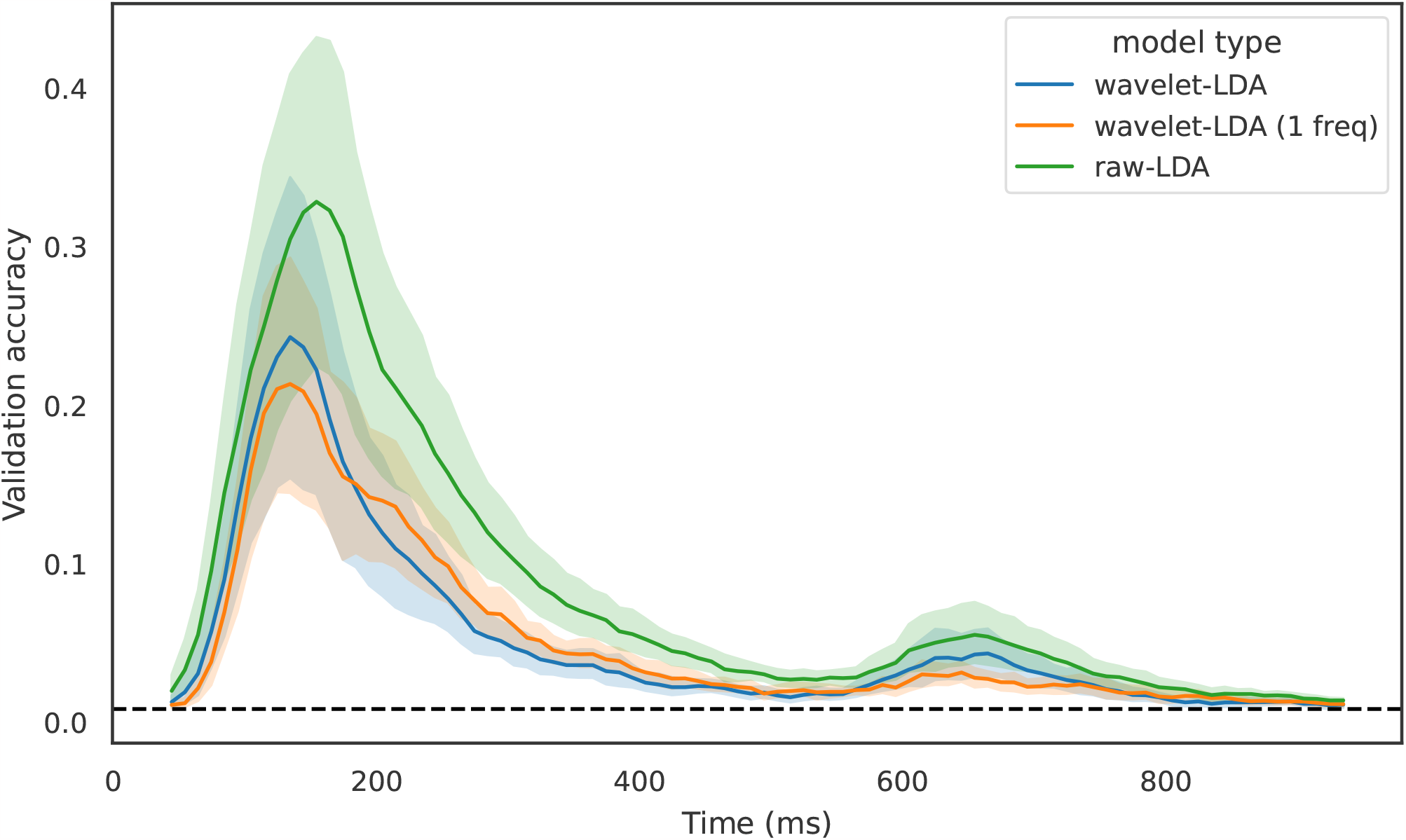
Comparison of our sliding window LDA-NN approach with LDA-NN using wavelet features on the 118-image dataset. The wavelet features are computed after the dimensionality reduction, with the same settings as in Higgins et al. (2022b). wavelet-LDA corresponds to using a concatenation of all frequency bands for training the LDA model, and wavelet-LDA (1 freq) uses a single frequency band (10Hz). Results are averaged across subjects, and shading indicates the 95% confidence interval across subjects.

## Footnotes

1 http://userpage.fu-berlin.de/rmcichy/fusion_project_page/main.html

