## Supplementary figures and images for "Interpretable full-epoch multiclass decoding for M/EEG"

### Inline Supplementary Video 1

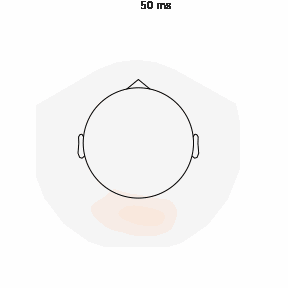
